# Long-Term Exposure to Elevated Lipoprotein(a) Levels, Parental Lifespan and Risk of Mortality

**DOI:** 10.1101/615898

**Authors:** Benoit J. Arsenault, William Pelletier, Yannick Kaiser, Nicolas Perrot, Christian Couture, Kay-Tee Khaw, Nicholas J. Wareham, Yohan Bossé, Philippe Pibarot, Erik S.G. Stroes, Patrick Mathieu, Sébastien Thériault, S. Matthijs Boekholdt

## Abstract

**Background:** Elevated Lipoprotein(a) (Lp[a]) levels are associated with a broad range of atherosclerotic cardiovascular diseases (CVD). The impact of high Lp(a) levels on human longevity is however controversial. Our objectives were to determine whether genetically-determined Lp(a) levels are associated with parental lifespan and to assess the association between measured and genetically-determined Lp(a) levels and long-term all-cause and cardiovascular mortality.

**Methods:** We determined the association between a genetic risk score of 26 single nucleotide polymorphisms weighted for their impact on Lp(a) levels (wGRS) and parental lifespan (at least one long-lived parent; father still alive and older than 90 or father’s age of death ≥90 or mother still alive and older than 93 or mother’s age of death ≥93) in 139,362 participants from the UK Biobank. A total of 17,686 participants were considered as having high parental lifespan. We also investigated the association between Lp(a) levels and all-cause and cardiovascular mortality in 18,720 participants from the EPIC-Norfolk study.

**Results:** In the UK Biobank, increases in the wGRS (weighted for a 50 mg/dL increase in Lp(a) levels) were inversely associated with a high parental lifespan (odds ratio=0.92, 95% confidence interval [CI]=0.89-0.94, p=2.7×10^−8^). During the 20-year follow-up of the EPIC-Norfolk study, 5686 participants died (2412 from CVD-related causes). Compared to participants with Lp(a) levels <50 mg/dL, those with Lp(a) levels ≥50 mg/dL had an increased hazard ratio (HR) for all-cause (HR=1.17, 95% CI=1.08-1.27) and cardiovascular (HR=1.54, 95% CI=1.37-1.72) mortality. Compared to individuals with Lp(a) levels below the 50^th^ percentile of the Lp(a) distribution (in whom event rates were 29.8% and 11.3%, respectively for all-cause and cardiovascular mortality), those with Lp(a) levels equal or above the 95^th^ percentile of the population distribution (≥70 mg/dL) had HRs of 1.22 (95% CI=1.09-1.37, event rate 37.5%) and 1.71 (95% CI=1.46-2.00, event rate 20.0%), for all-cause mortality and cardiovascular mortality, respectively.

**Conclusions:** Results of this study suggest a potentially causal effect of Lp(a) on human longevity, support the use of parental lifespan as a tool to study the genetic determinants of human longevity, and provide a rationale for a trial of Lp(a)-lowering therapy in individuals with high Lp(a) levels.

## INTRODUCTION

Lipoprotein(a) (Lp[a]) consists of a low-density lipoprotein (LDL) attached to apolipoprotein(a) (apo[a]) by a disulfide bond. Plasma levels of Lp(a) are predominantly explained by genetic variations at the *LPA* locus on chromosome 6. Approximately one in five individuals worldwide has an Lp(a) level (equal or above 50 mg/dL or 120 nmol/L) placing them at higher risk of a broad range of atherosclerotic cardiovascular disease (CVD) and calcific aortic valve stenosis.^1–5^ Despite the strong association between elevated Lp(a) levels and CVD risk, the evidence linking Lp(a) levels and Lp(a)-raising genetic variants with the risk of all-cause mortality is not as consistent. A 1998 study of healthy centenarians initiated a debate about the potential association between Lp(a) and longevity following their report that up to a quarter of healthy centenarians had high Lp(a) levels in absence of any atherosclerotic CVD.^6^ Another study of patients with documented coronary heart disease found no evidence of an association between high Lp(a) levels and all-cause mortality.^7^ This research question is of particular relevance as Lp(a)-lowering therapies are currently being developed and one of them (an antisense oligonucleotide against *LPA* called AKCEA-APO[a]-L_rx_)^8^ is expected to be tested in a large phase three cardiovascular outcomes trial (CVOT). Determining the association between high Lp(a) levels in large, prospective studies would inform on the potential of these therapies to extend lifespan in individuals with high Lp(a) levels.

The definition of what constitutes longevity in human genetic studies is highly debated and the lack of a universally recognized definition increases the possibility of biases and hinders external validation efforts, especially for case-control studies.^9^ The selection of appropriate controls is also almost as important as the selection of cases in such genetic association studies. Results of many studies on centenarians or long-lived individuals might have been confounded by the use of different birth cohorts of centenarians and controls, selection bias or survival bias. Parental lifespan is a novel and innovative tool that is increasingly used to study the genetic makeup of human longevity that considerably reduces selection bias as both cases and controls are uniformly recruited. Two recent genome-wide association studies identified variants at the *LPA* locus to be associated with shorter lifespan (as estimated by parental lifespan).^10,11^

The association between measured and genetically-determined Lp(a) levels and human longevity is controversial and despite evidence suggesting that *LPA* might be a locus influencing longevity, it is unknown if a concentration-dependent effect of Lp(a) levels on human longevity exists. In this study, we used a 2-sample Mendelian randomization (2SMR) study design to determine whether genetic variants associated with elevated Lp(a) levels are causally associated with human longevity, as estimated by parental lifespan, in the UK Biobank. We also investigated the association between measured and genetically-determined Lp(a) levels and long-term all-cause and cardiovascular mortality in another cohort from the United Kingdom, the European Prospective Investigation into Cancer and Nutrition (EPIC)-Norfolk study.

## METHODS

### Study populations

The association between *LPA* variants and parental lifespan was assessed in the UK Biobank, which includes approximately 500,000 individuals between 40 and 69 years old recruited between 2006 and 2010 in several centers in the United Kingdom.^12^ These analyses were conducted under UK Biobank data application number 25205. The association between genetically determined and measured Lp(a) levels and long-term all-cause and cardiovascular mortality was assessed in the EPIC-Norfolk study, which is a population-based study of 25,663 men and women aged between 45 and 79 years residing in Norfolk, UK. Participants were recruited by mail from age-sex registers of general practices in Norfolk. The design, methods of the study and baseline characteristics of the study participants have been described previously.^4,13^ At the baseline survey conducted between 1993 and 1997, participants completed a detailed health and lifestyle questionnaire. Lp(a) levels were measured with an immune-turbidimetric assay using polyclonal antibodies directed against epitopes in apolipoprotein(a) (Denka Seiken, Coventry, United Kingdom), as previously described.^14^ This assay has been shown to be apolipoprotein(a) isoform-independent.

### Outcomes ascertainment and definitions

Participants were asked the current age of their parents or the age at which their parents had died. We used the definition of Pilling et al.^15^ to define high parental lifespan in participants of the UK Biobank. Only participants between 55 and 69 years were included. Participants who were adopted, had missing information on age of parents’ death or who had parents who died at a young age (<46 for the father and <57 for the mother) were excluded from these analyses. Parents were separated into three categories: long-lived (father still alive and older than 90 or father’s age of death ≥90 and mother still alive and older than 93 or mother’s age of death ≥93), medium lived (age of death ≥66 and <89 for the father and ≥73 and <92 for the mother) and short lived (age of death ≥46 and <65 for the father and ≥57 and <72 for the mother). We defined high parental lifespan as at least one long-lived parent (i.e. long/long or long/medium). Analyses were also performed for high paternal lifespan (with maternal lifespan being long or medium) and for high maternal lifespan (with paternal lifespan being long or medium). A second, more stringent, outcome was defined as at least one parent with exceptional longevity (top 1% survival, i.e. age of death ≥95 for the father or ≥98 for the mother with the other parent either long- or medium-lived). The control group included participants with parents considered as short- or medium-lived (i.e. short/short, short/medium or medium/medium). Participants discordant for mothers’ and fathers’ age of death (one long-lived parent and one short-lived parent) were also excluded from the present analyses.

In EPIC-Norfolk, all individuals were flagged for mortality at the UK Office of National Statistics, with vital status ascertained for the entire cohort. Death certificates for all decedents were coded by trained nosologists according to the International Classification of Diseases (ICD) 9^th^ revision. In addition, participants admitted to hospital were identified by their unique National Health Service number by data linkage with ENCORE (East Norfolk Health Authority database), which identifies all hospital contacts throughout England and Wales for Norfolk residents. In EPIC-Norfolk, among 18,720 individuals with Lp(a) measurement, 5686 died (2412 from CVD) during the follow-up.

### Genotyping and selection of genetic instruments

Samples were genotyped with the Affymetrix UK BiLEVE Axiom array or the Affymetrix UK Biobank Axiom Array. On the UK Biobank, phasing and imputation were performed centrally using the Haplotype Reference Consortium (HRC) reference panel.^16^ Samples with call rate <95%, outlier heterozygosity rate, gender mismatch, non-white British ancestry, related samples (second degree or closer), samples with excess third-degree relatives (>10), or not used for relatedness calculation were excluded. Burgess et al.^17^ recently used a genetic risk score (GRS) of 43 single nucleotide polymorphisms (SNP) that explained approximately 60% of the variance in Lp(a) levels in four large cohorts (R^2^ measure of linkage disequilibrium <0.4). The marginal effect of these SNPs on Lp(a) levels was obtained. To derive an estimation of genetically-determined Lp(a) levels, we included 26 SNPs from the report of Burgess et al.^17^ that had a minor allele frequency equal or above 0.005. We also only included independent SNPs (R^2^<0.2). In EPIC-Norfolk, we used two SNPs that had the strongest impact on Lp(a) levels (rs10455872 and rs3798220) in the study of Clarke et al.^18^

### Statistical analyses

To evaluate the association between genetically-determined Lp(a) levels and parental lifespan in the UK Biobank, we performed 2SMR, in which the effect of the selected SNPs on Lp(a) levels were obtained from Burgess et al.^17^ and the effect of the SNPs on parental lifespan were assessed in the UK Biobank. First, we separated individuals in the UK Biobank into quartiles based on wGRS distribution and performed logistic regression adjusting for age, sex and the first 10 ancestry-based principal components to document the association between genetically-elevated Lp(a) and parental lifespan. Second, we obtained effect estimates (adjusted for the minor allele frequency of each variant) by 50 mg/dL increase in Lp(a), a threshold recently reported by Langsted et al.^19^ We performed inverse-variance weighted (IVW)-MR by performing a meta-analysis of each Wald ratio (the effect of the genetic instrument on Lp[a] levels divided by its effect on parental lifespan). To determine the significance of the associations, a bootstrap method was used. A 2-tailed p-value was calculated using a z-test from 100,000 random simulations. IVW-MR is considered as one of the simplest ways to obtain MR estimates using multiple SNPs. The limitation of IVW-MR is the assumption that SNPs do not have pleiotropic effects (effects on other variables than the trait of interest). To determine the presence of unmeasured pleiotropy, we performed MR-Egger in which a nonzero y-intercept is allowed in order to assess violation of IVW-MR, as described by Bowden et al.^20^ These analyses were performed using R (Version 3.5.1). In EPIC-Norfolk, Cox regression was used to calculate hazard ratios and corresponding 95% confidence interval for the risk of all-cause and cardiovascular mortality associated with various thresholds of measured Lp(a) levels and two SNPs associated with high Lp(a) levels. Hazard rations for all-cause and cardiovascular mortality were obtained before and after adjusting for cardiovascular risk factors (age, sex, smoking, body mass index, systolic blood pressure, diabetes mellitus and creatinine) when evaluating measured Lp[a] levels and age and sex when evaluating Lp[a]-raising SNPs). We estimated the difference in survival between those with high (equal or above the 95^th^ percentile) versus low Lp(a) levels (below the 50^th^ percentile) in age-equivalent terms by dividing the beta coefficient for all-cause mortality associated with high versus low Lp(a) levels by the beta coefficient difference in all-cause mortality associated with one year increase in age, as previously described.^21,22^ These analyses were performed using SPSS software (Version 12.0.1, Chicago, IL).

## RESULTS

### Genetically-elevated Lipoprotein(a) and parental lifespan in the UK Biobank

Of the 139,362 UK Biobank participants included in this analysis, 17,686 were considered as having high parental lifespan and 2932 were defined as having one parent with exceptional longevity (top 1% survival). In the sex-specific analyses investigating paternal and maternal survival, 8976 individuals were considered as having high paternal lifespan and 10,137 were considered as having high maternal lifespan. Regardless of how longevity was defined, genetically-determined Lp(a) (whether examined as quartiles of the wGRS or as continuous GRS) was negatively associated with a high parental lifespan in the UK Biobank. The odds ratios for a high parental lifespan per 50 mg/dL increase in Lp(a) as well as in the study population separated into quartiles of the Lp(a) wGRS are presented in Figure 1.

**Figure 1.**
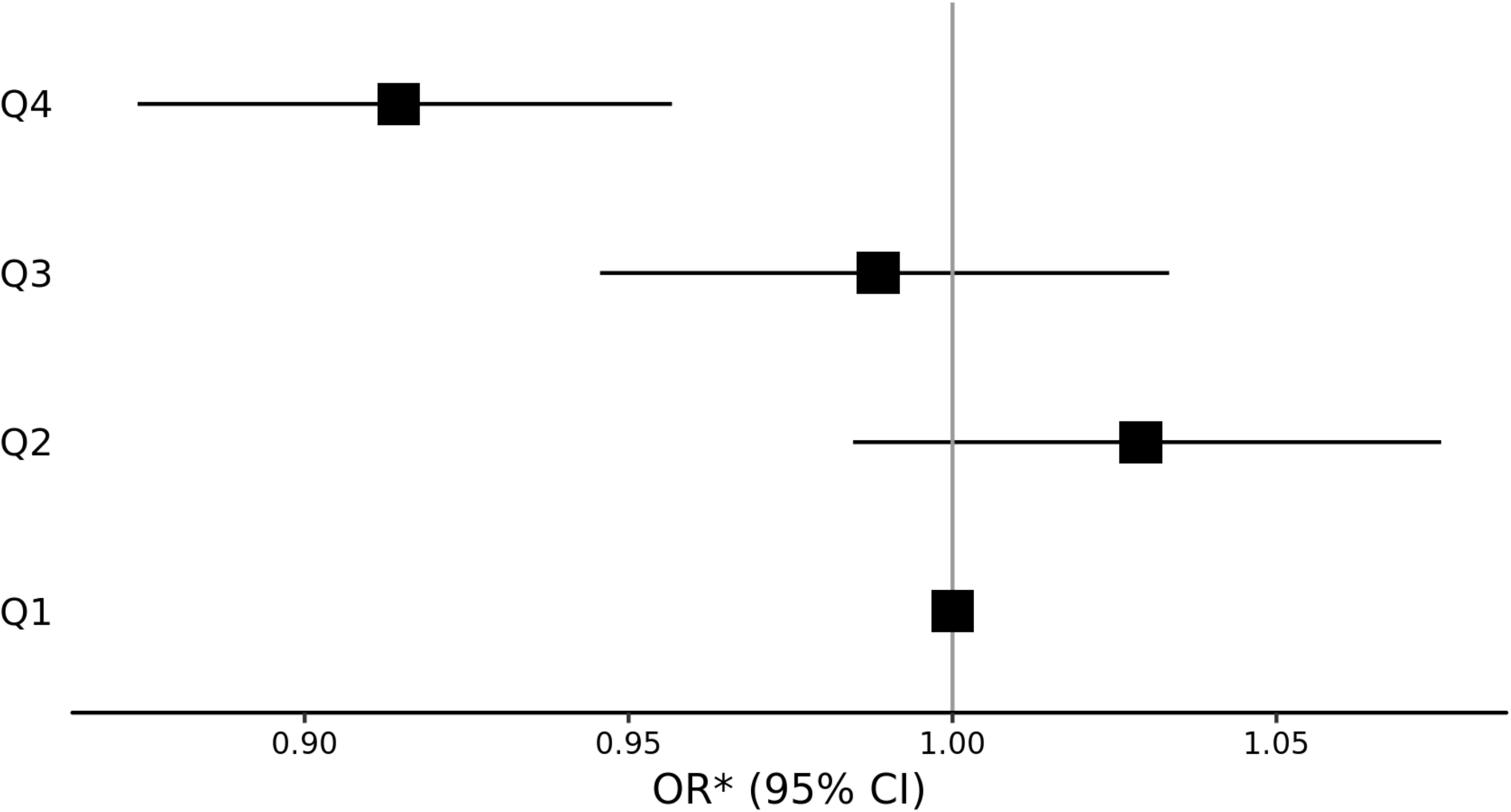
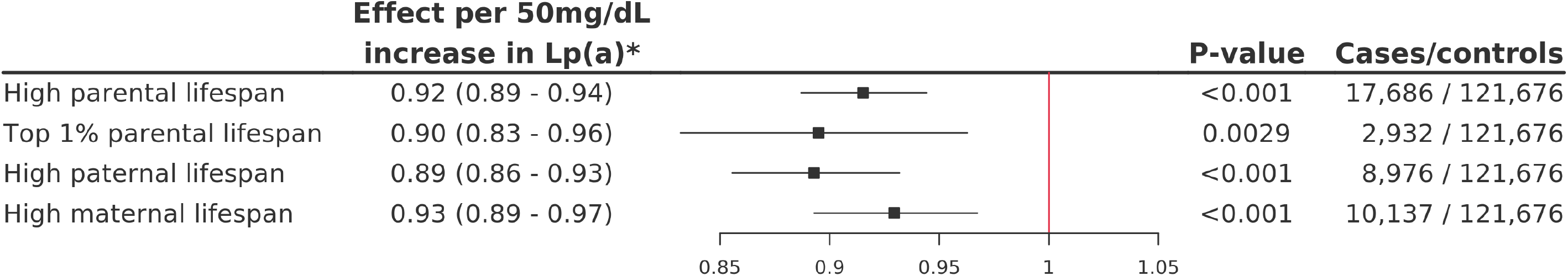
Impact of the *LPA* genetic instruments on higher parental lifespan. A) Odds ratio (OR) and 95% confidence interval (CI) for high parental lifespan in participants of the UK Biobank separated into quartiles of the *LPA* weighted genetic risk score (wGRS). B) OR and 95% CI for high parental lifespan, top 1% parental lifespan, high paternal lifespan and high maternal lifespan associated with a 50 mg/dL increase in the *LPA* wGRS in the UK Biobank. *Adjusted for age, sex and the 10 first ancestry-based principal components.

Figure 2 presents the association between the effects of the 26 *LPA* SNPs on Lp(a) levels and high parental lifespan. Higher genetically-determined Lp(a) levels were associated with lower chances of having high parental lifespan. We obtained estimates of causal effects of Lp(a) levels on parental lifespan in the UK Biobank using IVW-MR and Egger-MR (Table 1). Egger-MR analysis revealed that there was no evidence of horizontal pleiotropy in the two outcomes that combined paternal and maternal lifespan. There was however evidence of horizontal pleiotropy when maternal lifespan only was investigated (P-value of intercept=0.04).

**Table 1.** Estimates of causal effects of lipoprotein(a) levels on parental lifespan in the UK Biobank.

|  | IVW-MR |  | Egger-MR |  |  |
| --- | --- | --- | --- | --- | --- |
|  | Slope estimate<br>(SD) | p-value | Slope estimate<br>(SD) | p-value | Intercept (p-value)* |
| High parental lifespan | -0.0020 (0.0004) | $3.17 \times 10^{-9}$ | -0.0026 (0.0004) | $1.68 \times 10^{-8}$ | 0.0038 (0.08) |
| Top 1% parental lifespan | -0.0027 (0.0008) | $1.97 \times 10^{-5}$ | -0.0020 (0.0011) | 0.06 | -0.0048 (0.35) |
| High paternal lifespan | -0.0027 (0.0005) | $1.43 \times 10^{-8}$ | -0.0028 (0.0005) | $1.18 \times 10^{-5}$ | 0.0006 (0.85) |
| High maternal lifespan | -0.0017 (0.0004) | $1.60 \times 10^{-4}$ | -0.0025 (0.0006) | $4.00 \times 10^{-5}$ | 0.0056 (0.04) |

IVW-MR indicates inverse-variance weighted Mendelian randomization and SD indicates standard deviation. \*A p-value <0.05 indicates that the y-intercept of the Mendelian randomization regression line is significantly different from 0, suggesting unbalanced pleiotropy.

**Figure 2.**
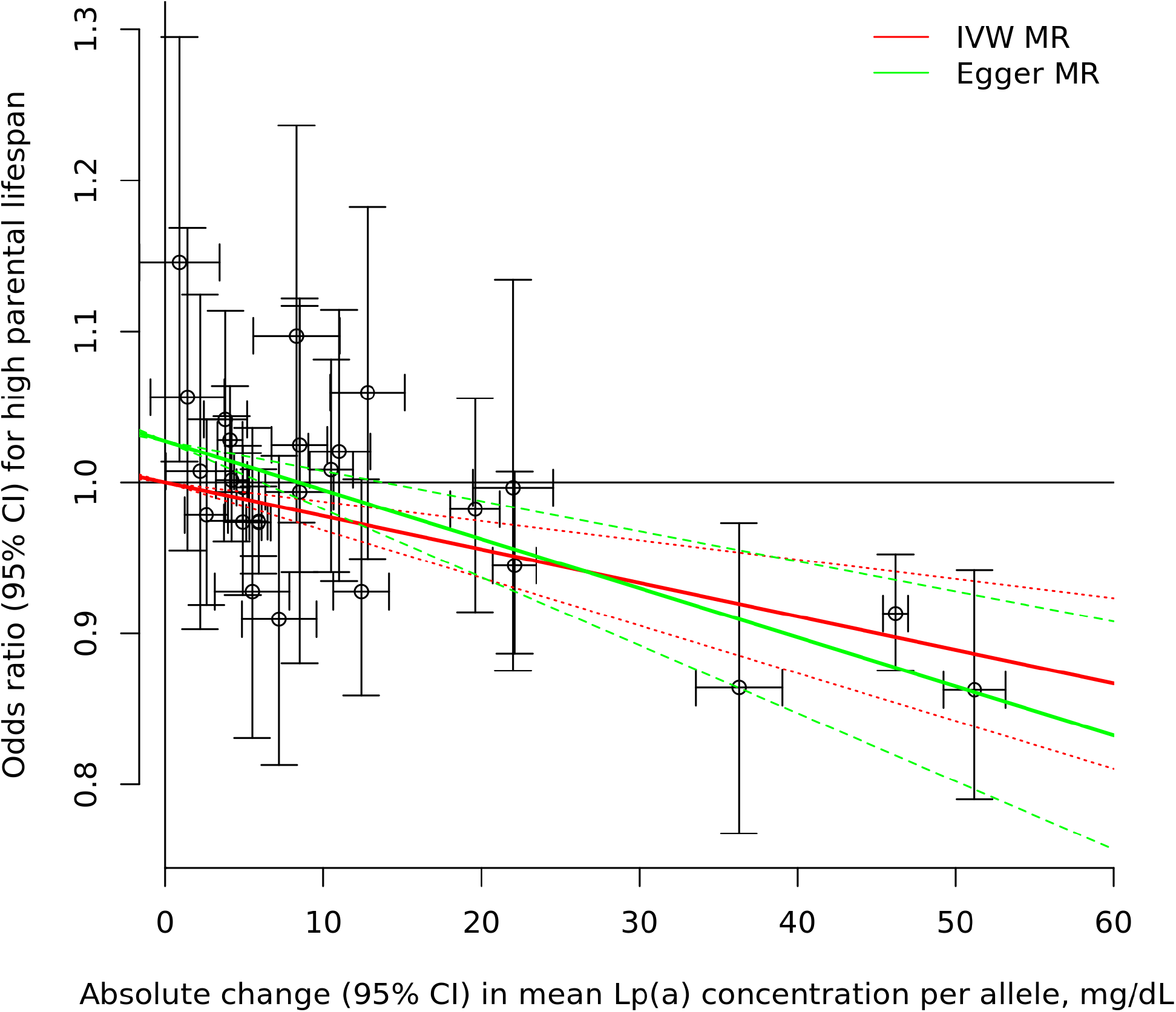
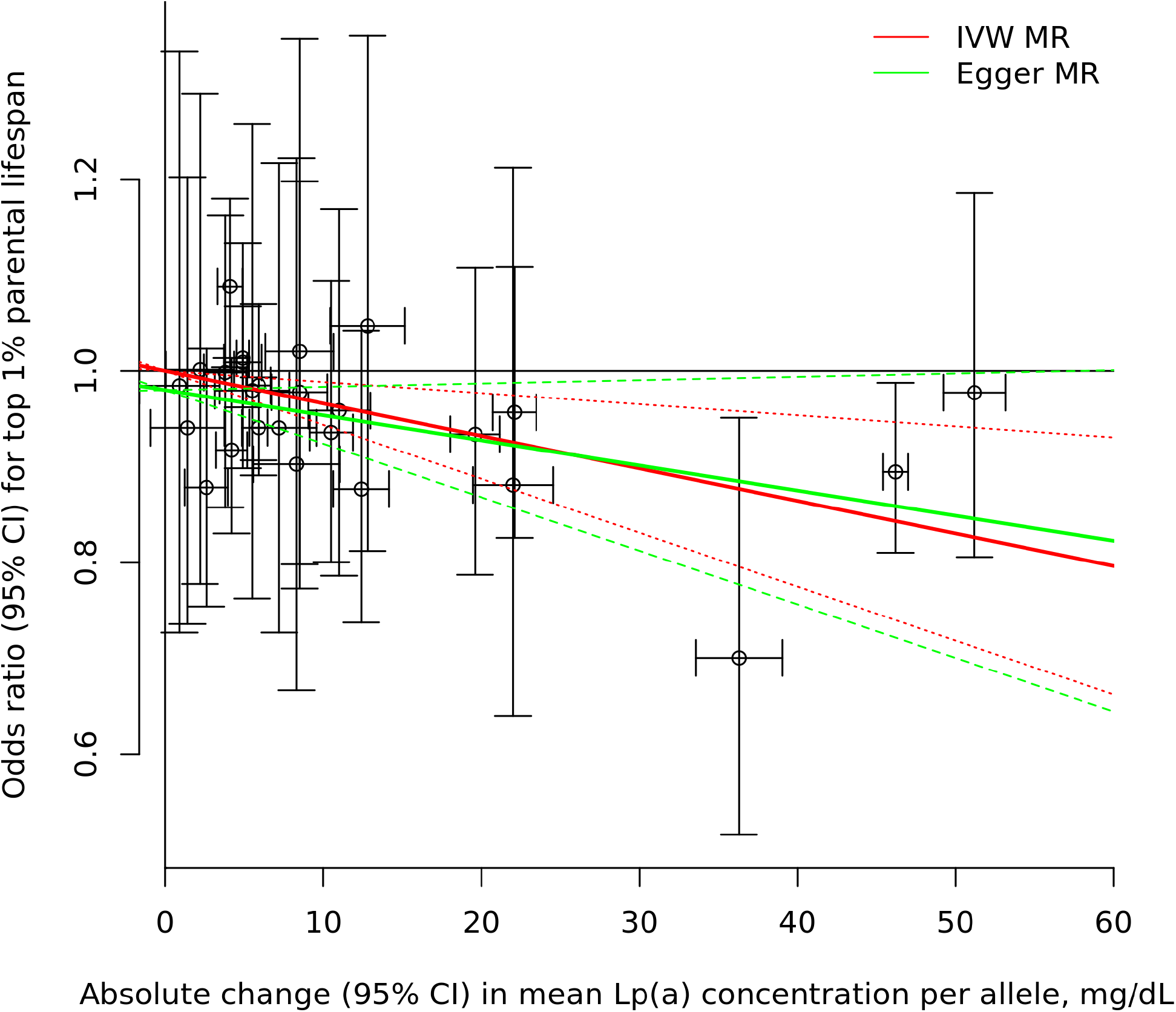
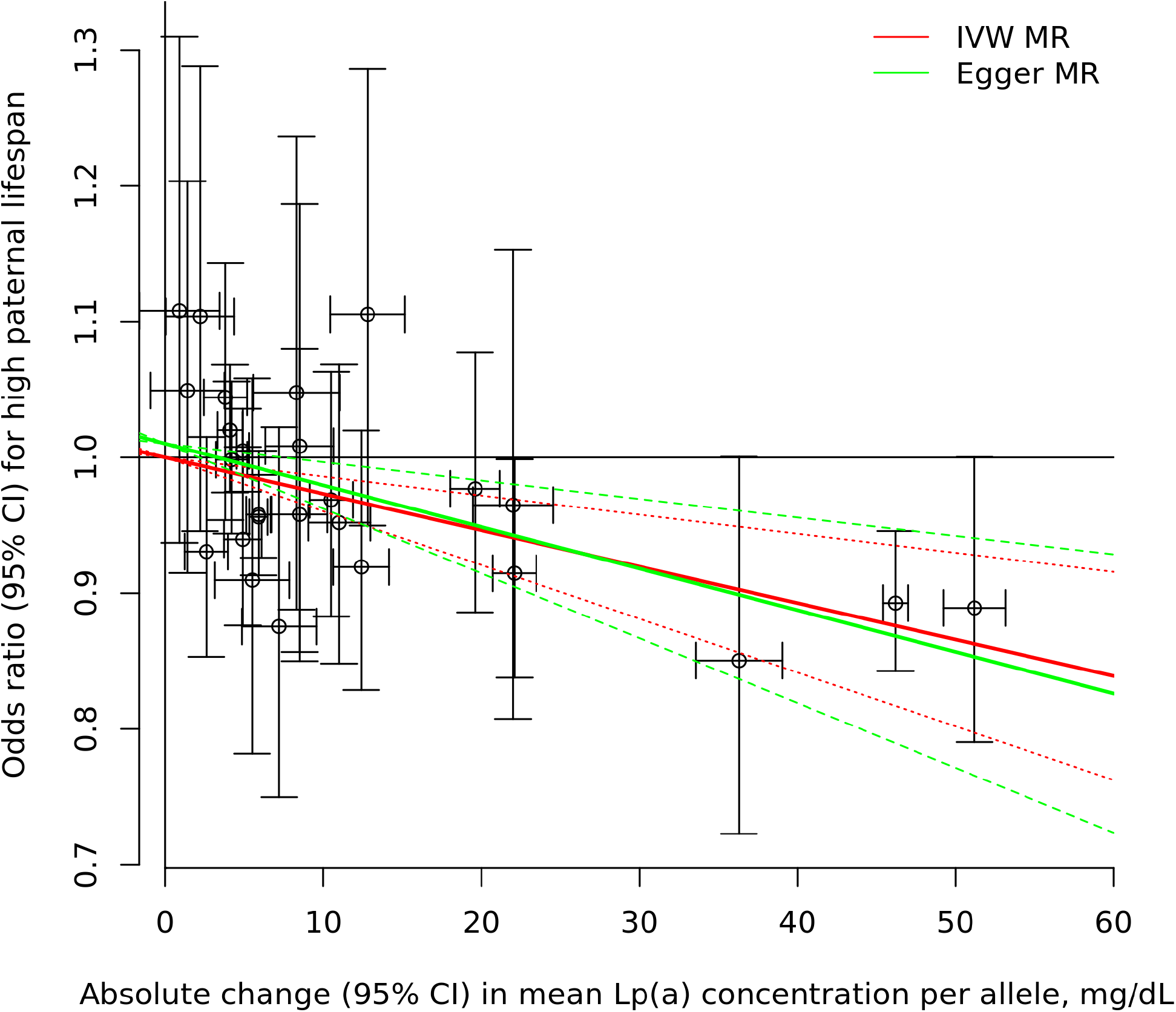
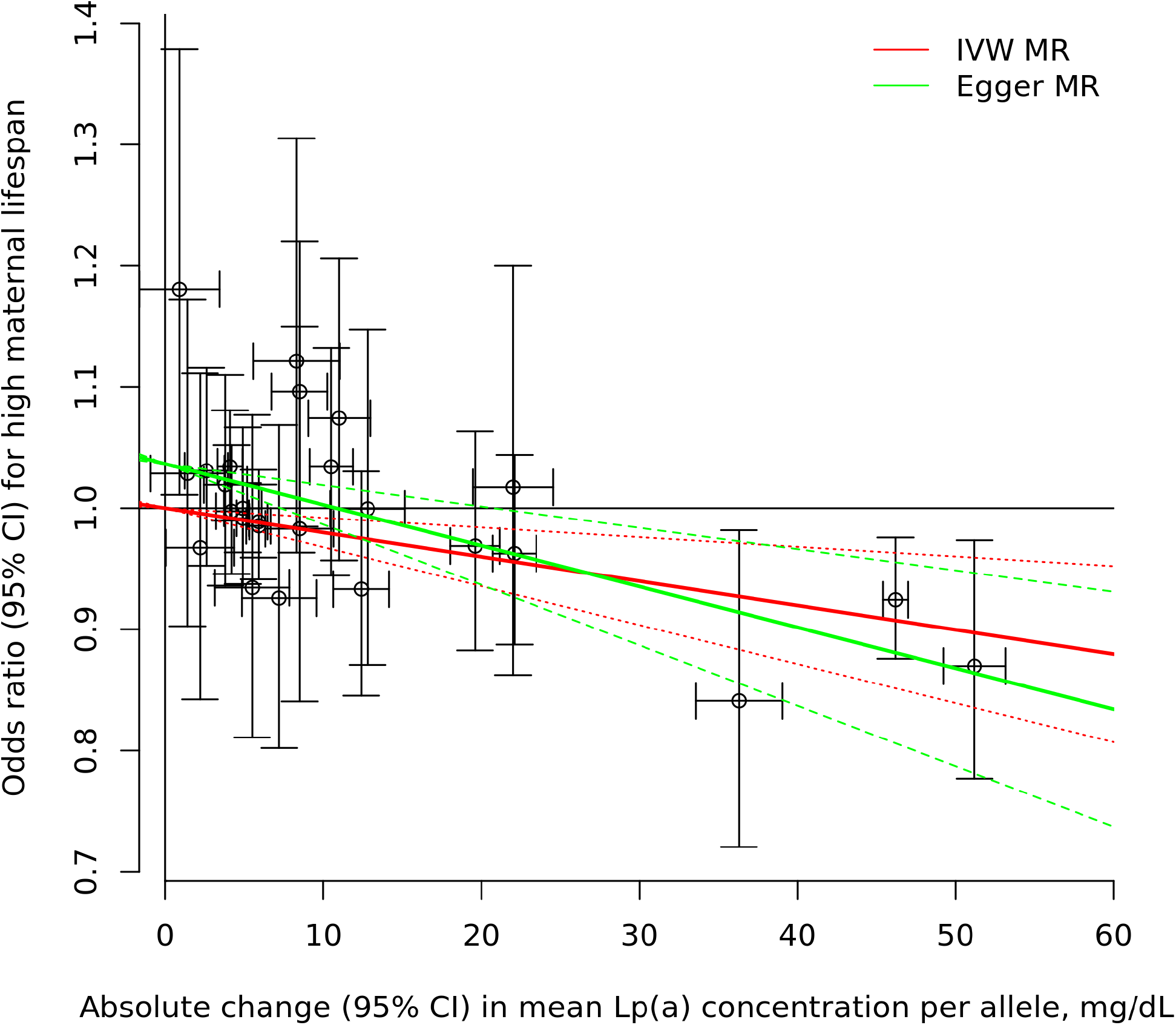
Association between SNPs at the *LPA* locus and higher parental lifespan [A], top 1% parental lifespan [B], high paternal lifespan [C] and high maternal lifespan [D]) in the UK Biobank. Each plotted point represents the effect of a single genetic variant on lipoprotein(a) levels (x-axis) and a high parental lifespan (y-axis). The red line represents the regression slope using the inverse-variance weighted method and the green line represents the regression slope using the Egger method. IVW-MR indicates inverse-variance weighted Mendelian randomization.

### Measured and genetically-elevated lipoprotein(a) and mortality in EPIC-Norfolk

The baseline characteristics of EPIC-Norfolk study participants by Lp(a) levels are presented in Table 2. Participants of the EPIC-Norfolk study were followed for an average of 20 years. Compared to participants with Lp(a) levels <50 mg/dL, those with Lp(a) levels ≥50 mg/dL had an increased hazard ratio (HR) of both all-cause and cardiovascular mortality (Figure 3). In sex-specific analyses, the association of high Lp(a) levels with cardiovascular mortality was observed in both men and women while the association of high Lp(a) levels with all-cause mortality was only statistically significant in men. No associations were found with the risk of non-cardiovascular mortality in the entire group and in the sex-specific analyses.

**Table 2.**
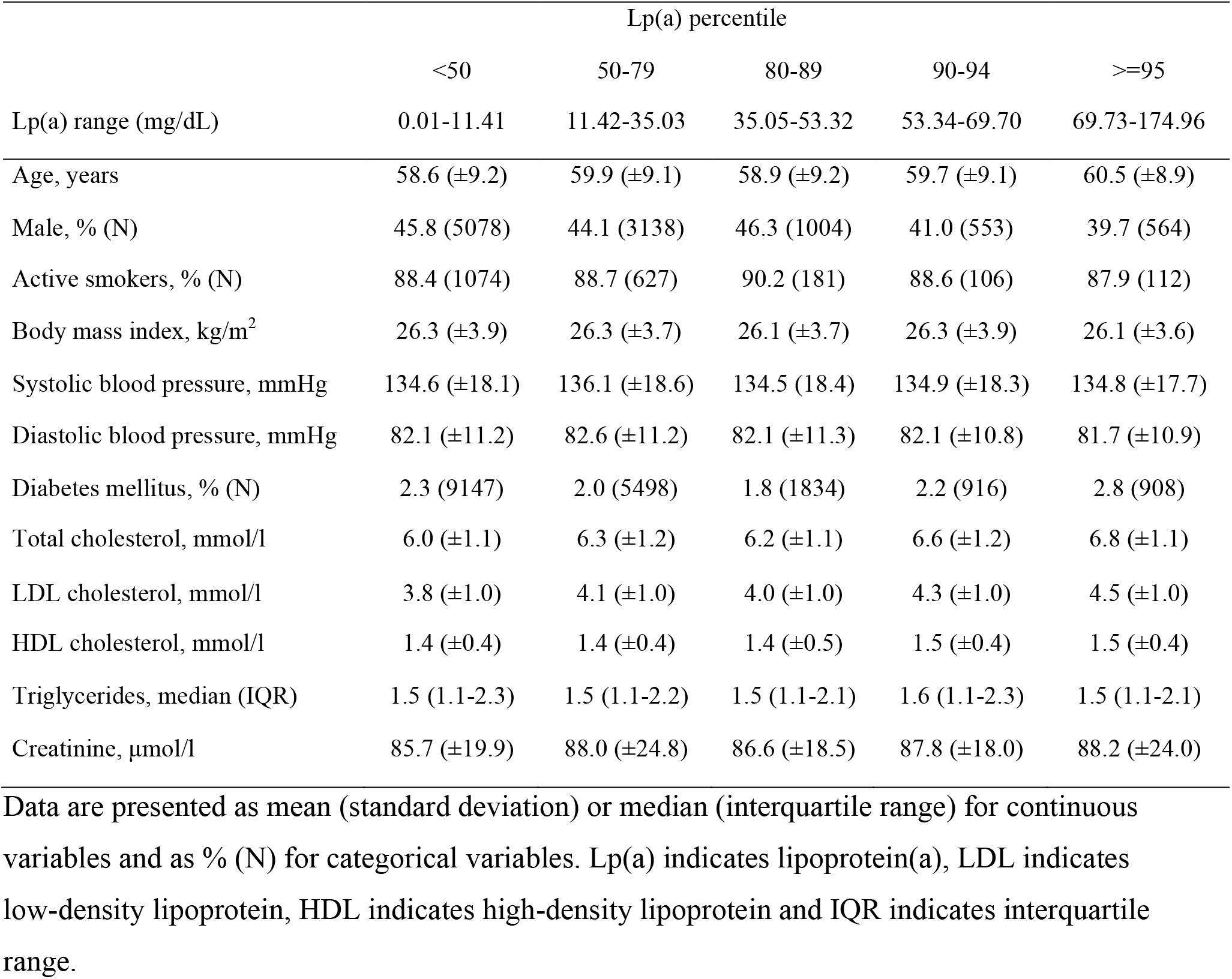
Baseline clinical characteristics of the EPIC-Norfolk study population and the study population by Lipoprotein(a) levels percentiles.

|  | Lp(a) percentile |  |  |  |  |
| --- | --- | --- | --- | --- | --- |
|  | <50 | 50-79 | 80-89 | 90-94 | ≥95 |
| Lp(a) range (mg/dL) | 0.01-11.41 | 11.42-35.03 | 35.05-53.32 | 53.34-69.70 | 69.73-174.96 |
| Age, years | 58.6 (±9.2) | 59.9 (±9.1) | 58.9 (±9.2) | 59.7 (±9.1) | 60.5 (±8.9) |
| Male, % (N) | 45.8 (5078) | 44.1 (3138) | 46.3 (1004) | 41.0 (553) | 39.7 (564) |
| Active smokers, % (N) | 88.4 (1074) | 88.7 (627) | 90.2 (181) | 88.6 (106) | 87.9 (112) |
| Body mass index, kg/m<sup>2</sup> | 26.3 (±3.9) | 26.3 (±3.7) | 26.1 (±3.7) | 26.3 (±3.9) | 26.1 (±3.6) |
| Systolic blood pressure, mmHg | 134.6 (±18.1) | 136.1 (±18.6) | 134.5 (18.4) | 134.9 (±18.3) | 134.8 (±17.7) |
| Diastolic blood pressure, mmHg | 82.1 (±11.2) | 82.6 (±11.2) | 82.1 (±11.3) | 82.1 (±10.8) | 81.7 (±10.9) |
| Diabetes mellitus, % (N) | 2.3 (9147) | 2.0 (5498) | 1.8 (1834) | 2.2 (916) | 2.8 (908) |
| Total cholesterol, mmol/l | 6.0 (±1.1) | 6.3 (±1.2) | 6.2 (±1.1) | 6.6 (±1.2) | 6.8 (±1.1) |
| LDL cholesterol, mmol/l | 3.8 (±1.0) | 4.1 (±1.0) | 4.0 (±1.0) | 4.3 (±1.0) | 4.5 (±1.0) |
| HDL cholesterol, mmol/l | 1.4 (±0.4) | 1.4 (±0.4) | 1.4 (±0.5) | 1.5 (±0.4) | 1.5 (±0.4) |
| Triglycerides, median (IQR) | 1.5 (1.1-2.3) | 1.5 (1.1-2.2) | 1.5 (1.1-2.1) | 1.6 (1.1-2.3) | 1.5 (1.1-2.1) |
| Creatinine, µmol/l | 85.7 (±19.9) | 88.0 (±24.8) | 86.6 (±18.5) | 87.8 (±18.0) | 88.2 (±24.0) |

Data are presented as mean (standard deviation) or median (interquartile range) for continuous variables and as % (N) for categorical variables. Lp(a) indicates lipoprotein(a), LDL indicates low-density lipoprotein, HDL indicates high-density lipoprotein and IQR indicates interquartile range.

**Figure 3.**
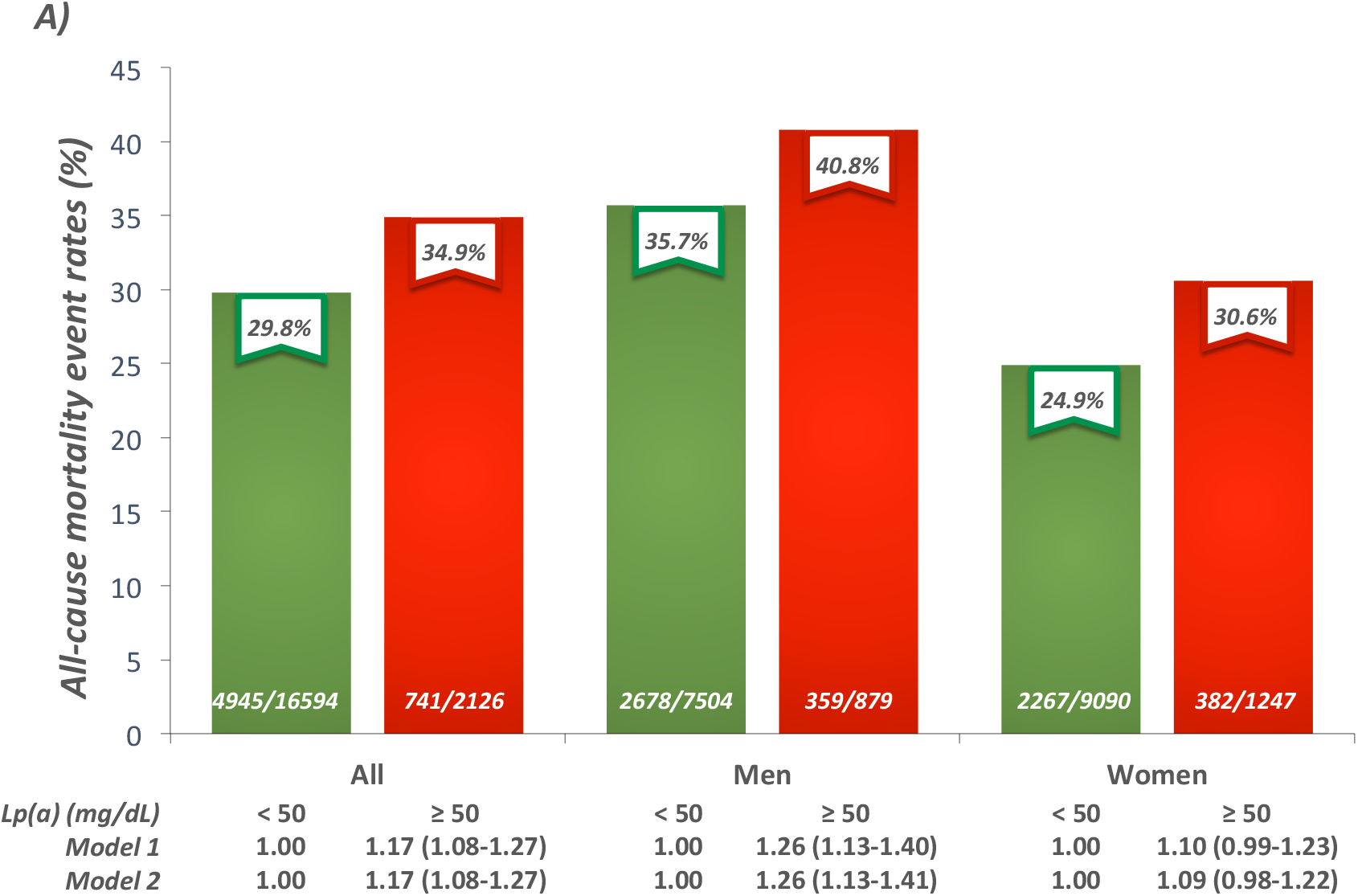
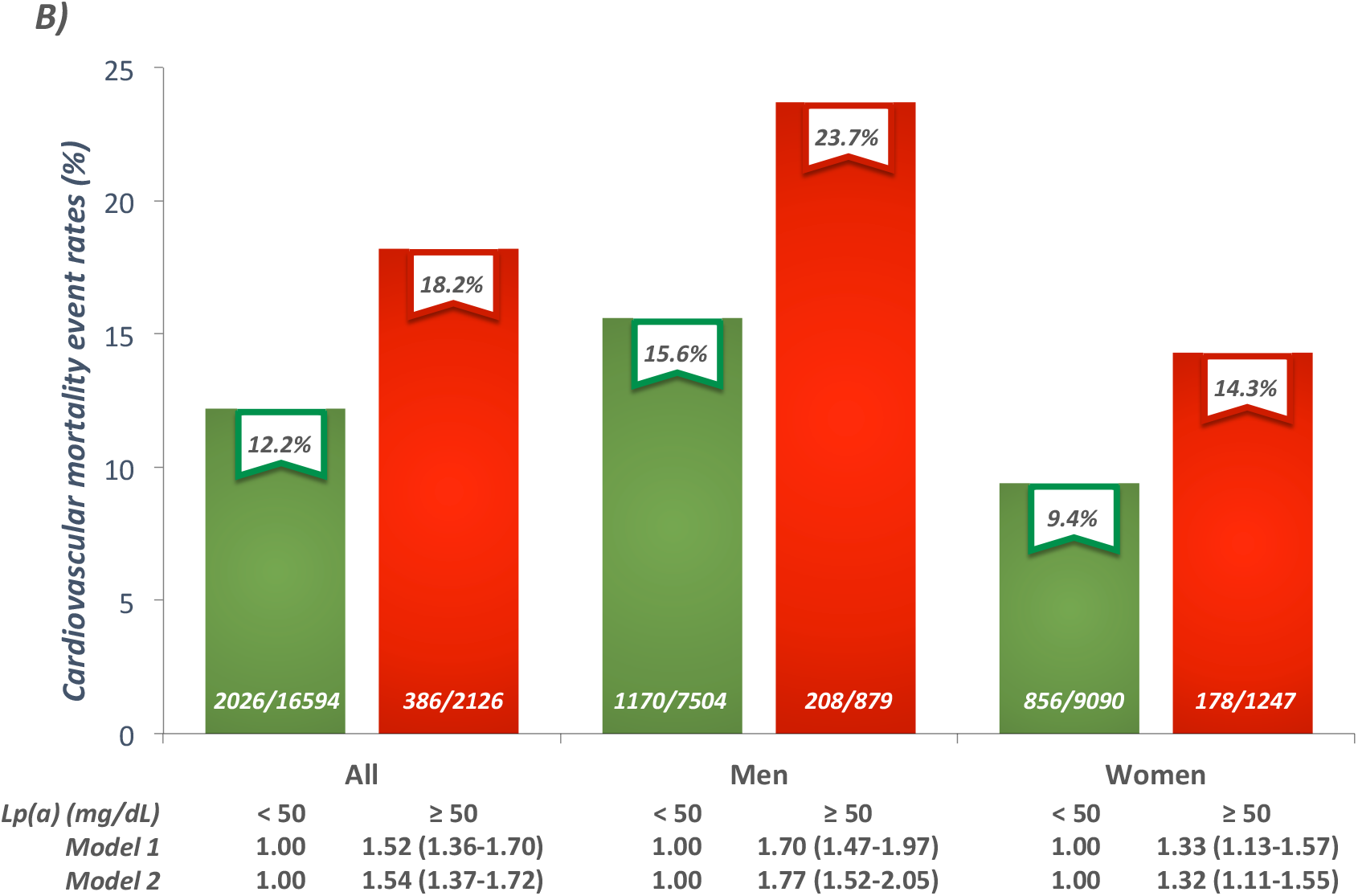
Event rates and hazard ratios for all-cause (A) and cardiovascular (B) mortality in participants of the EPIC-Norfolk study with low or high lipoprotein(a) levels (< or ≥50 mg/dL), as well as in men and women separately. Model 1 is adjusted for age and sex. Model 2 is adjusted for age, sex, smoking, body mass index, systolic blood pressure, diabetes mellitus and estimated glomerular filtration rate.

Next, we sought to determine whether there was a concentration-dependent effect of Lp(a) on all-cause and cardiovascular mortality in participants above the 50^th^ percentile of the Lp(a) distribution. Results presented in Figure 4 suggest that the risk of all-cause mortality becomes statistically significant in participants above the 90^th^ percentile of the Lp(a) distribution while the risk of cardiovascular mortality becomes statistically significant in participants above the 80^th^ percentile of the Lp(a) distribution. The relative risks for all-cause and cardiovascular mortality were highest in participants with Lp(a) levels equal or above the 95^th^ percentile.

**Figure 4.**
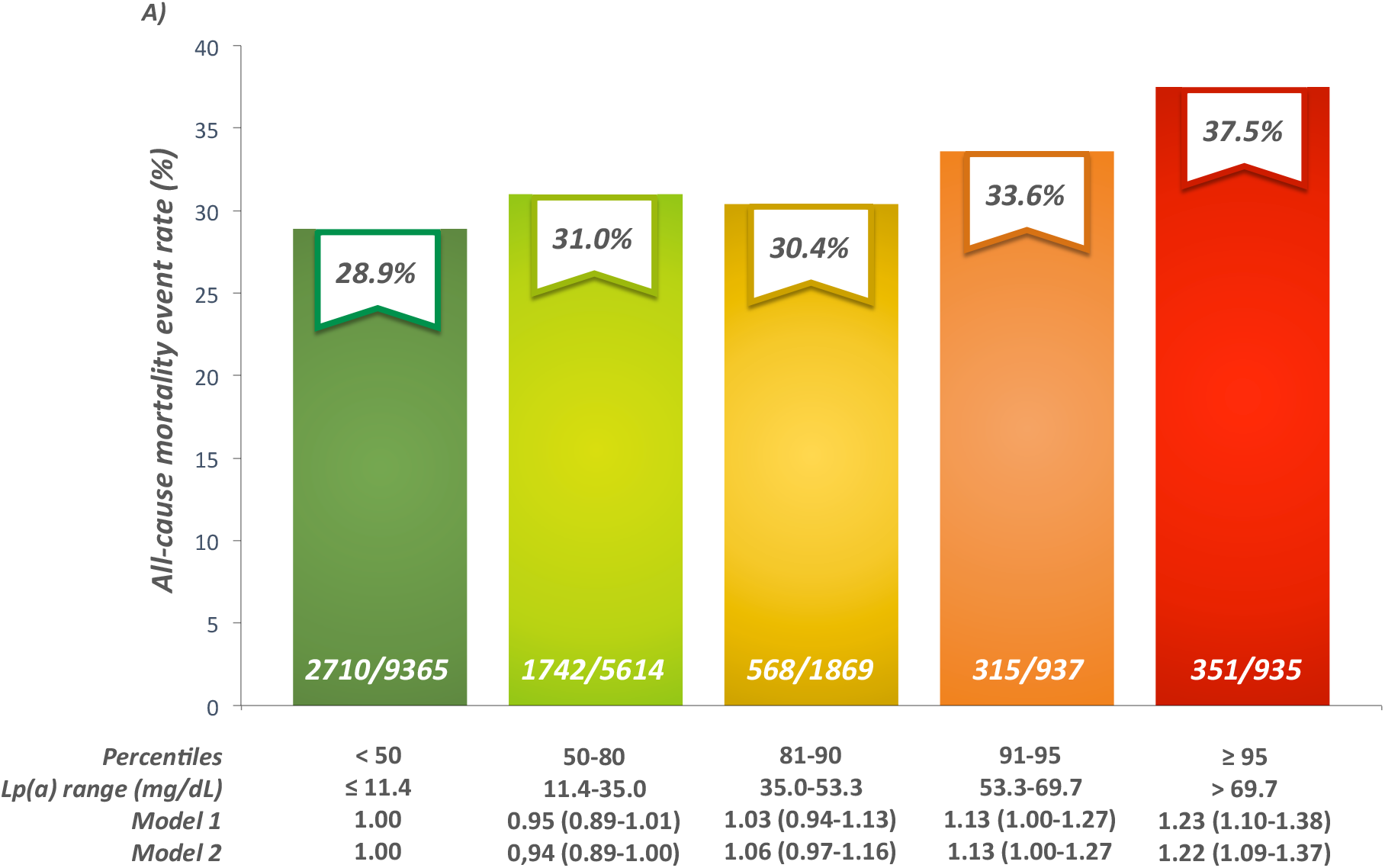
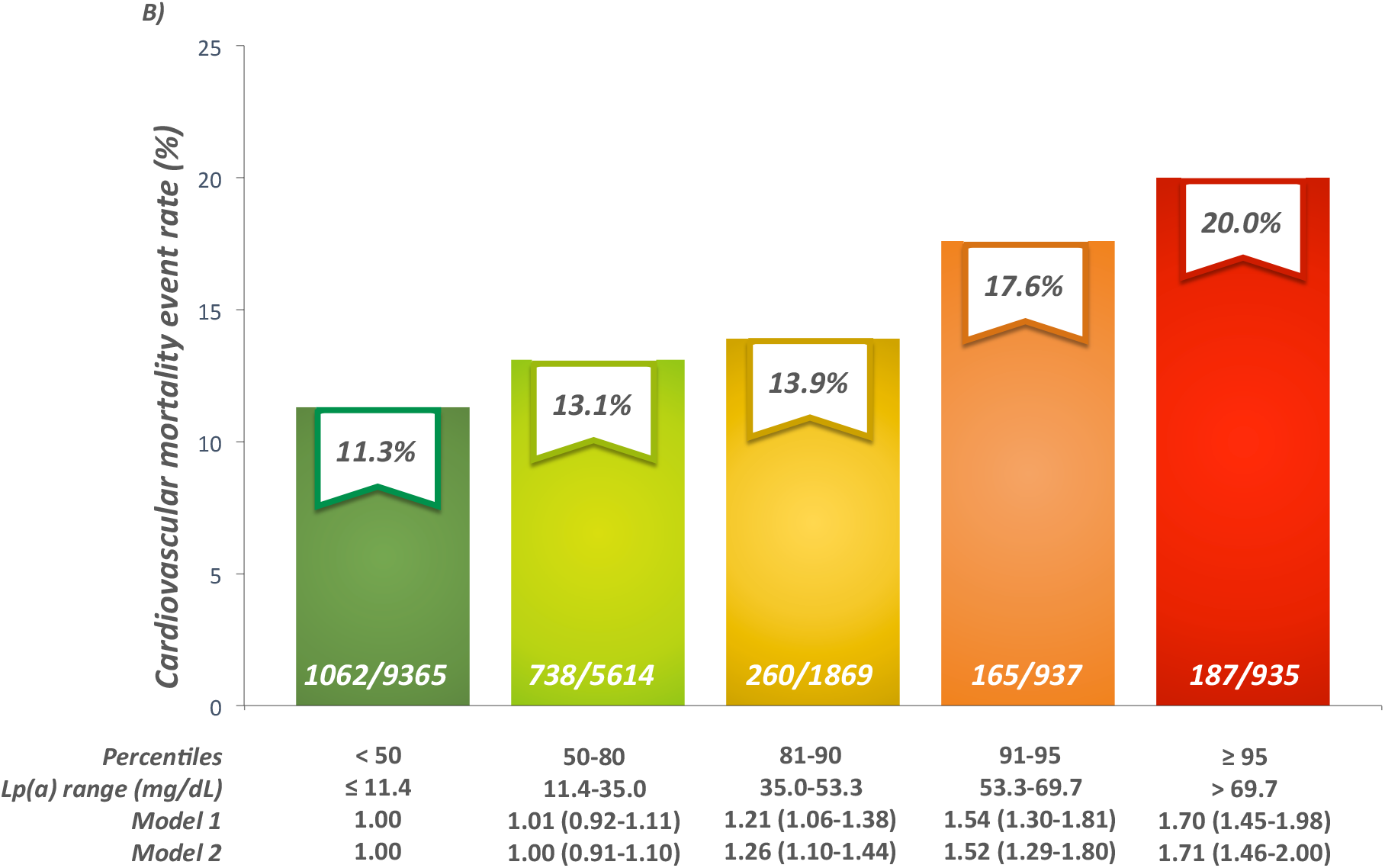
Event rates and hazard ratios for all-cause (A) and cardiovascular (B) mortality in participants of the EPIC-Norfolk study by extreme high lipoprotein(a) levels. Model 1 is adjusted for age and sex. Model 2 is adjusted for age, sex, smoking, body mass index, systolic blood pressure, diabetes mellitus and estimated glomerular filtration rate.

From the Cox model, the beta coefficient for all-cause mortality associated with each year increase in chronological age was 0.127 (± standard error 0.003). The beta coefficient for a comparison between high (equal or above the 95^th^ percentile) versus low Lp(a) levels (below the 50^th^ percentile) was 0.194 (± standard error 0.064), which is equivalent to approximately 1.5 years in chronological age for all-cause mortality risk. This analysis suggests that the mortality risk for those with Lp(a) levels equal or above the 95^th^ percentile is equivalent to being 1.5 years older in chronological age.

Next, we investigated the association between two SNPs with a strong effect on Lp(a) levels (rs10455872 and rs3798220) and all-cause and cardiovascular mortality. For rs10455872, compared to non-carriers (AA genotype; event rate of 29.6% and 12.2%, respectively for all-cause and cardiovascular mortality), those who carried at least one Lp(a) raising allele (AG or GG genotype) were at higher risk for all-cause (HR=1.14 [95% CI, 1.07-1.22], event rate of 31.9%) and cardiovascular mortality (HR=1.23 [95% CI, 1.11-1.36], event rate of 13.9%). For rs3798220, compared to non-carriers (TT genotype; event rate of 29.9% and 12.4%, respectively for all-cause and cardiovascular mortality), those who carried at least one Lp(a) raising allele (TC or CC genotype) were however not at significantly higher risk for all-cause (HR=1.03 [95% CI, 0.90 −1.18], event rate of 29.3%) and cardiovascular mortality (HR=1.16 [95% CI, 0.94 −1.42, event rate of 13.4%). Compared to individuals without Lp(a)-raising allele, those with only one Lp(a)-raising allele (in rs10455872 or rs3798220) had an increased risk of both all-cause and cardiovascular mortality (Figure 5). Those with two or more Lp(a)-raising alleles had an even higher risk of all-cause mortality and cardiovascular mortality, although the association with cardiovascular mortality did not reach statistical significance (HR=1.40, 95% CI=0.98-2.00). However, there were only 202 individuals in that subcategory, including 30 who died of CVD. No associations were found with the risk of non-cardiovascular mortality.

**Figure 5.**
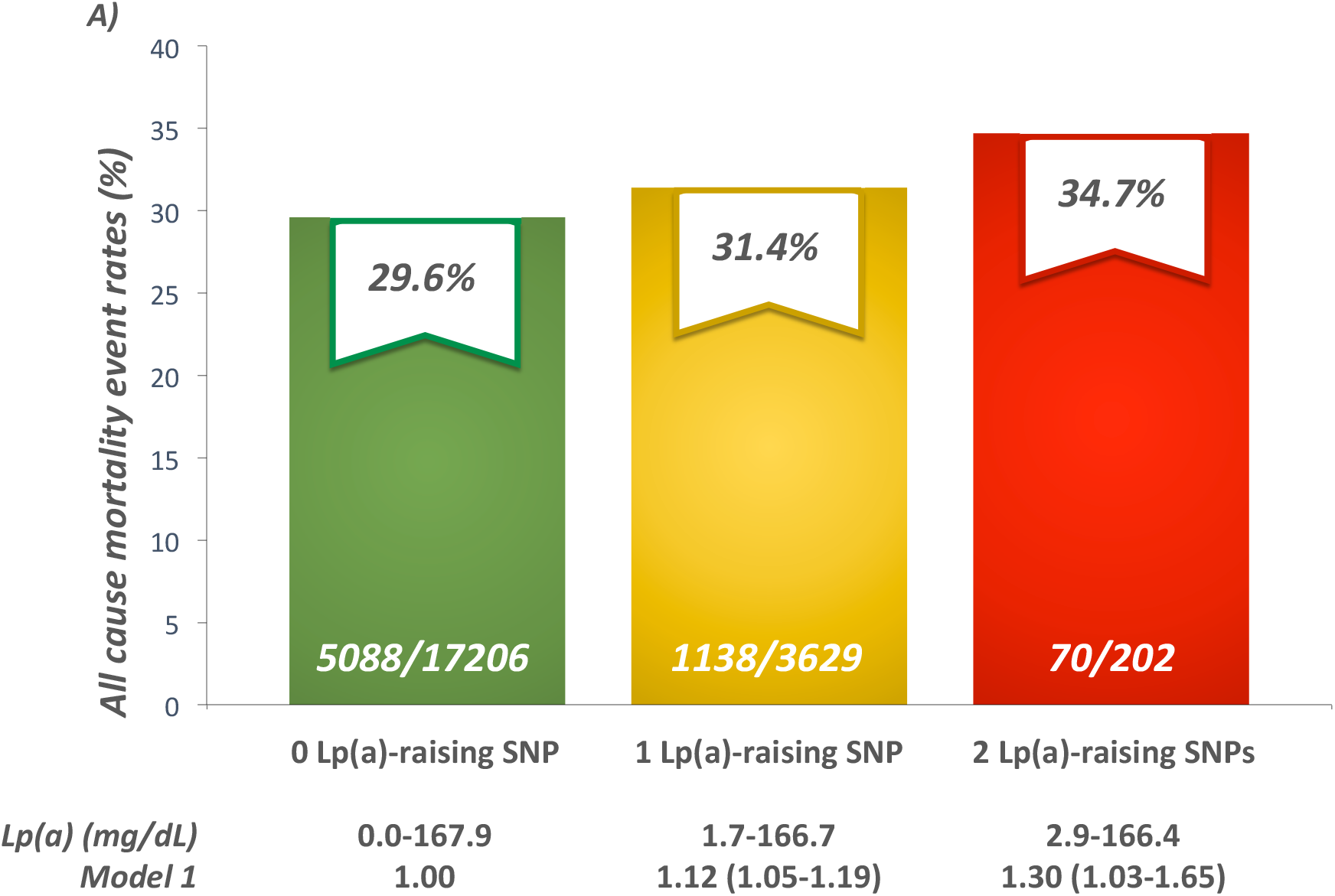
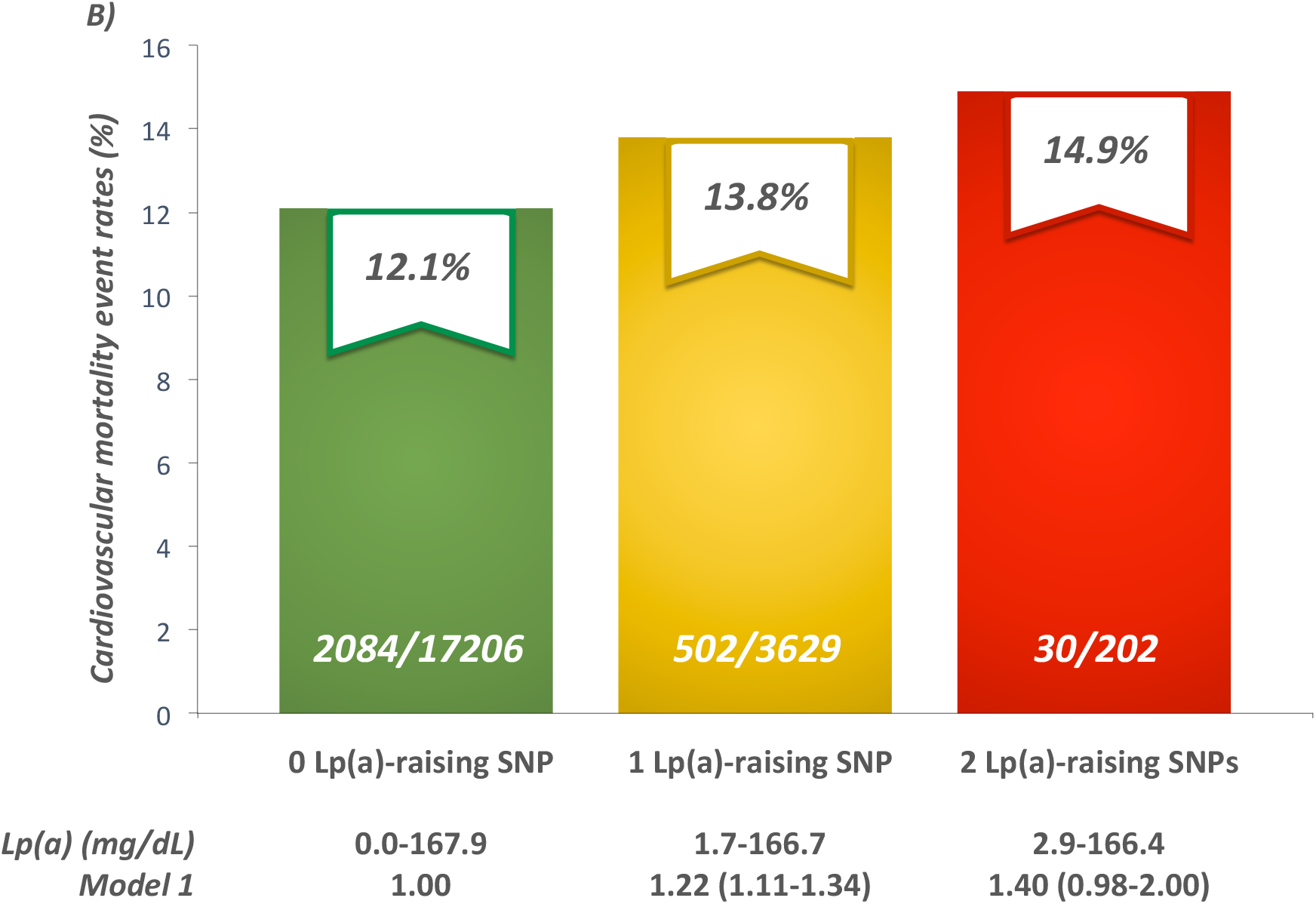
Event rates and hazard ratios for all-cause (A) and cardiovascular (B) mortality in participants of the EPIC-Norfolk study by number of lipoprotein(a)-raising alleles. Model 1 is adjusted for age and sex.

## DISCUSSION

Results of our MR study suggest that genetically-determined Lp(a) levels are causally associated with parental lifespan in participants of the UK Biobank. We also provide evidence that genetically-determined as well as measured Lp(a) levels are associated with the long-term risk of all-cause and cardiovascular mortality in 18,720 participants of the EPIC-Norfolk prospective population study followed for an average of 20 years where the mortality risk for those with Lp(a) levels equal or above the 95^th^ percentile was equivalent to being 1.5 years older in chronological age. Altogether, our results suggest that variants at *LPA*, through an increase in absolute Lp(a) levels, are important determinants of human longevity (Figure 6).

**Figure 6.**
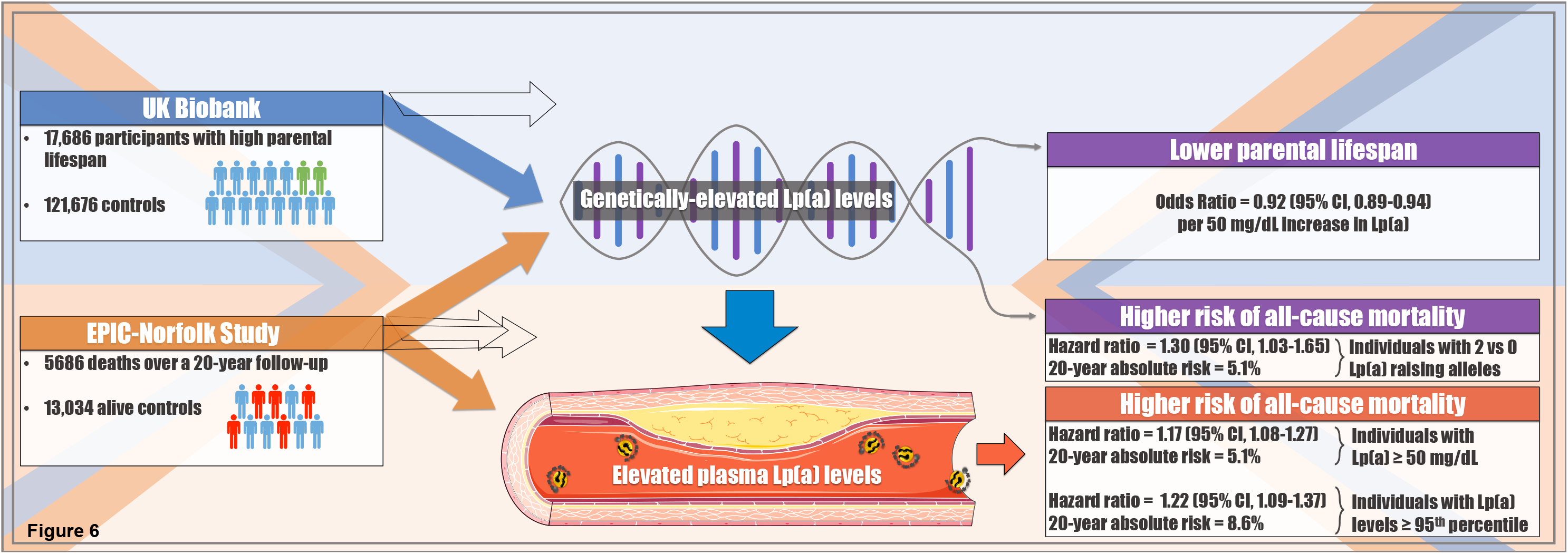
Long-term exposure to elevated lipoprotein(a) levels is associated with shorter parental lifespan and long-term risk of mortality.

In a 2017 genetic association study that sought to identify variants associated with parental lifespan, Joshi et al.^11^ identified four loci including the *LPA* locus to be associated with parental lifespan at the genome-wide significance level. In a follow-up study of over a million parental lifespans, Timmers et al.^10^ confirmed the association between variants in *LPA* and parental lifespan. Interesting results were also recently reported by Zenin et al.^23^ who have shown that variants in *LPA* may be associated with disease-free survival (also known as healthspan) in the UK Biobank, thereby suggesting that lower Lp(a) might not only be associated with longer lifespan, but also with healthy living into old age. These studies however did not investigate the potential effect of genetically-elevated Lp(a) levels on parental lifespan or healthspan using robust genetic analyses such as MR. By reporting a significant effect of high Lp(a) levels on shorter parental lifespan using MR, our study strengthens the case for Lp(a) as a causal determinant of human longevity.

Results of our study also provide validation for the use of parental lifespan for the study of the genetic determinants of human longevity. The association between our trait of interest and parental lifespan here reported using a 2SMR study design and subsequent validation in a long-term prospective study that included 18,720 apparently healthy individuals with 5686 incident mortality cases, also support the use of MR as an innovative tool or surrogate to study the genetic makeup of human longevity. MR studies could be useful to determine whether suspected biological determinants of longevity have a causal role in the etiology of this complex trait.

A previous study of healthy centenarians showed that almost a quarter of healthy centenarians had high Lp(a) levels despite little to no evidence of atherosclerotic CVD.^6^ Limitations of that study included the very small sample size (only 75 centenarians), the measurement of Lp(a) with an assay that may not have considered apo(a) isoform size and the possibility of survival bias. Because they are less prone to reverse-causality, prospective studies are better suited than case-control studies to assess the long-term effect of risk factors such as Lp(a) on cardiometabolic outcomes and all-cause mortality. In 2009, the Emerging Risk Factor Collaboration reported a positive association between high Lp(a) levels and all-cause mortality in a meta-analysis of 24 long-term prospective studies.^5^ More recently, investigators of two Danish prospective population studies (Copenhagen City Heart Study [CCHS] and Copenhagen General Population Study [CGPS]) also suggested a possible association between high levels of Lp(a) and all-cause and cardiovascular mortality in the general population.^19^ In the Danish studies, compared to participants in the bottom 50^th^ percentile of the Lp(a) distribution (all-cause mortality event rate of 14.2% and cardiovascular mortality event rate of 3.6%), those with Lp(a) above the 95^th^ percentile had a hazard ratio for all-cause mortality of 1.20 (95% CI, 1.10-1.30, event rate of 16.5%) and a hazard ratio for cardiovascular mortality of 1.50 (95% CI, 1.28-1.76, event rate of 5.0%). In our study, using comparable subgroups, we found that the hazard ratio for all-cause and cardiovascular mortality were remarkably consistent with the Danish studies. In our study however, the absolute risk of all-cause and cardiovascular mortality in participants with Lp(a) levels above the 95^th^ percentile was 8.6% and 8.7% higher than the group with low Lp(a) levels, respectively for all-cause and cardiovascular mortality. The absolute risk associated with high Lp(a) reported here is considerably higher than what was observed in the CCHS and CGPS (2.5% and 1.4%, respectively for all-cause and cardiovascular mortality), most likely reflecting the longer follow-up period in EPIC-Norfolk. However, in contrast with the CCHS and CGPS that reported a null association between the Lp(a)-raising variant rs10455872 and all-cause and cardiovascular mortality, we found a strong dose-response association between the number of rs10455872-G alleles and all-cause and cardiovascular mortality, thereby confirming that absolute Lp(a) levels are strong predictors of all-cause and cardiovascular mortality.

In the UK Biobank, we found that the effect of SNPs on Lp(a) levels inversely correlated with the odds of high paternal and maternal lifespan. In EPIC-Norfolk, we found that Lp(a) appeared to be more strongly associated with long-term all-cause and cardiovascular mortality risk in men compared to women. Additional studies will be needed to document potential sex-differences in the association between high Lp(a) and mortality.

We investigated the association between high Lp(a) and all-cause and cardiovascular mortality using a well-accepted cut-off value of 50 mg/dL in the EPIC-Norfolk study. Our results suggest that compared to participants with Lp(a) levels below that threshold, those with Lp(a) levels equal or above 50 mg/dL had a 5.1% increase in absolute event rate and a 17% (95 % CI, 8-27%) increase in the hazard ratio for all-cause mortality and a 6.0% increase in absolute event rate and a 54% (95 % CI, 37-72%) increase in the hazard ratio for cardiovascular mortality. It is unknown if lowering Lp(a) with investigative therapies will change the risk trajectory of individuals with high Lp(a) levels and bring it back to the level of risk of individuals with low Lp(a) levels. Although our results were obtained in a general population study, they provide the rationale for an Lp(a)-lowering trial in the prevention of total and cardiovascular mortality in high-risk patients with elevated Lp(a) levels. In conclusion, our results highlight Lp(a) as an important genetic determinant of human longevity. Whether lowering Lp(a) levels will ultimately prolong life and promote healthy living into old age in patients with high Lp(a) levels will need to be tested in long-term randomized clinical trials.

## ACKNOWLEDGEMENTS

We would like to thank all study participants. BJA and ST hold junior scholar awards from the *Fonds de recherche du Québec: Santé*. BJA is a consultant for Novartis and has received research funding from Pfizer, Amgen and Ionis Pharmaceuticals. The EPIC-Norfolk Study is funded by Cancer Research UK grant number 14136 and the Medical Research Council grant number G1000143. PP holds the Canada Research Chair in Valvular Heart Disease and his research program is supported by a Foundation Scheme Grant from CIHR. PM holds a FRQS Research Chair on the Pathobiology of Calcific Aortic Valve Disease and is a consultant for Casebia Therapeutics. YB holds a Canada Research Chair in Genomics of Heart and Lung Diseases.

